# Animal-Origin-Free Method for Generating Blood Vessel Organoids

**DOI:** 10.1101/2025.11.26.690667

**Authors:** Alexander Hoffmann, David Schorn, Jakob Thönig, Teodor E. Yordanov

## Abstract

**Background:** Blood vessel organoids (BVOs) represent a promising tool for modeling vascular diseases, drug screening, and regenerative therapies. However, current protocols for BVO generation are complex, labor-intensive, and reliant on animal-derived extracellular matrices (ECM) such as Matrigel, limiting reproducibility, scalability, and clinical applicability.

**Methods:** We developed a simplified, animal-origin-free protocol for BVO generation that addresses current limitations and enables high-throughput automated workflows. The method employs ultra-low attachment 96-well U-bottom plates for standardized aggregation and differentiation of human induced pluripotent stem cells (hiPSCs) in a human derived collagen-based extracellular matrix. Unlike conventional protocols where aggregates are embedded in a two-layer ECM, our approach utilizes a single-layer, which we termed “sitting drop”. This innovative approach requires considerably fewer materials and handling steps and is compatible with high-throughput automated machines.

**Results:** BVO generation utilizing the here described optimized protocol resulted in the formation of BVOs with reproducible morphology and cellular composition. Flow cytometry confirmed the presence of CD31⁺ endothelial cells and PDGFRβ⁺ pericytes in BVOs, generated in sitting drops, with cell population percentages comparable to those observed in traditional two-layer BVO cultures. *In vivo* transplantation of mature BVOs in a mouse full-thickness skin wound model demonstrated successful integration of BVO derived cells into host vessels, highlighting their potential in cell-based therapies.

**Conclusion:** Our study presents a robust and animal-origin-free method for BVO generation based on single-layer “sitting drop” cultures. This protocol maintains cellular integrity while enhancing reproducibility and automation-readiness, paving the way for high-throughput screening and clinical translation of vascular organoid technology.

**Highlights:**

- Entire blood vessel organoid (BVO) workflow performed in a single ultra-low attachment-96 plate
- Fully animal-origin-free: no Matrigel or Geltrex required throughout the protocol
- Robust generation of BVOs using human collagen-based ECM
- High-throughput compatible and automation-ready “sitting drop” culture system
- Integration of BVO-derived cells into host vessels *in vivo*

## 1. Introduction

Organoids are three-dimensional *in vitro* models of organs derived from stem cells, including those representing the small intestine (1), liver (2), lung (3), pancreas (4), brain (5), and blood vessels (6), among others. These models hold immense potential for both basic and translational research, enabling high-throughput drug screening, reducing reliance on animal experiments, and paving the way for innovative cell-based therapies and personalized medicine. In particular, Blood Vessel Organoids (BVOs) have emerged as a promising tool for studying vascular diseases such as diabetic vasculopathy (7) and atherosclerosis (8).

Despite their potential, current protocols for generating BVOs face key limitations. Most approaches are based on the protocol by Wimmer et al. (6), which involves the use of animal-derived extracellular matrices such as Matrigel. These matrices are not only expensive and non-compliant with good manufacturing practice (GMP), but also exhibit considerable batch-to-batch variability, significantly compromising reproducibility (9, 10). In addition, the conventional two-layer embedding technique requires manual excising, transferring organoids between plates, which makes the workflow labor-intensive, prone to errors and is poorly suited for automation.

To overcome these limitations, we developed a simplified and animal-origin-free protocol for BVO generation that is compatible with GMP. Our method enabled aggregate formation, differentiation and organoid formation in a single plate using standardized ultra-low attachment 96-well U-bottom plates (ULU-96) and a human-collagen-based extracellular matrix (HC-ECM). Instead of the conventional two-layer embedding method, we introduced a one-step extracellular matrix (ECM) embedding approach that simplifies handling and enhances high-throughput compatibility, which we refer to as the “sitting drop” method.

This work establishes a robust platform for reproducible, animal-origin-free BVO production, paving the way for translational applications in regenerative medicine and disease modeling.

## 2. Methods

### 2.1. Cell culture

Human Induced Pluripotent Stem Cells (hiPSC) were purchased from Pluristyx (201 Elliott Ave W, Seattle, WA 98119, USA). Cells were cultured on Nunc™ Cell-Culture Treated Multidishes 6 Well-plate (ThermoFischerScientific, 150675) precoated with 2ml DPBS, calcium, magnesium (DPBS +/+) (Gibco, 14040117) and 2.5 µg/ml Biolaminin 521 LN (Biolamina, LN521-05). Plates were incubated in a MCO-230AICUV IncuSafe CO2 Incubator (PHCEurope). Animal origin free culture media was prepared using 47.5 ml TeSR™-AOF Basal medium (STEMCELL, 100-0402) with 2.5 ml TeSR™-AOF 20X Supplement (STEMCELL, 100-0403) and 100µl Antibiotic Normocin (InvivoGen, NOL-44-09) and was used for up to 2 weeks stored at 4°C. For passaging, 70-80% confluent cells were washed with DPBS, no calcium, no magnesium (DPBS -/-) (Gibco, 14190144) before cell dissociation using ReLeSR (Stemcell, 100-0483) incubated for 1 minute at 37°C. ReLeSR was aspirated, and the plate was incubated for 5 minutes at 37°C with no medium. Cells were resuspend using 1 ml of culture medium and seeded on precoated plates with a minimum dilution of 1:10.

### 2.2. Protocol for Blood Vessel Organoid (BVO) Formation and Maturation in Single Layer Sitting Drops in 96 Well Plates

**Day 0: Aggregation Formation**

1. **Prepare Aggregation Formation Medium**

- Mix STEMdiff™ BVO Aggregation Basal Medium (STEMCELL, Cat. No.: 100-0652) with STEMdiff™ BVO Aggregation Supplement (STEMCELL, Cat. No.: 100-0653) in a 4:1 ratio (medium: supplement) at RT.
- After mixing at RT, warm the Aggregation Formation Medium to 37 °C.
2. **Cell collection and counting**

- Dilute 0.5 M EDTA (Invitrogen UltraPure™ 0.5 M EDTA, pH 8.0) to 0.5 mM in DPBS (50 µL EDTA + 50 mL DPBS/PBS without Mg²⁺ and Ca²⁺) and warm it in the water bath to 37 °C.
- Thaw Accutase (StemPro® Accutase® Cell Dissociation Reagent, Cat. No.: A1110501).
- Aspirate medium from the cells and add 0.6 mL of warm 0.5 mM EDTA in DPBS.
- Incubate at 37 °C and 5% CO₂ for 5 minutes.
- Carefully remove the plate from the incubator, ensuring not to shake.
- Aspirate EDTA solution from the cells without removing the cells.
- Add 0.6 mL of RT Accutase.
- Incubate at 37 °C and 5% CO₂ for 5 minutes.
- Check whether cell detachment is successful by gently tapping the plate.
- Once cells are detached, add 2 × 0.7 mL of Aggregate Formation Medium, collect the cells, and transfer the cell suspension to a sterile 15 mL conical tube.
- Rinse the well with 1 mL of Aggregate Formation Medium and add it to the conical tube.
- Use a P1000 tip to pipette up and down to ensure a homogeneous cell distribution before counting.
- Count the cells using trypan blue and a Countess system, a hemocytometer or a comparable system.
- Calculate the volume of cell suspension required to seed 1000 cells per well and transfer it to a new 15 mL conical tube.
3. **Centrifuge and resuspend cells**

- Centrifuge the cells at 250-300 × g for 3-5 minutes.
- Add ROCK inhibitor (Y-27632) (Miltenyi, Cat. No.: 130-104-169) to the Aggregation Formation Medium to achieve a final concentration of 50 µM.
- Warm medium to 37 °C before use.
- Remove the supernatant from the cell pellet.
- Resuspend the cell pellet in Aggregation Seeding Medium (with ROCK inhibitor Y-27632) to a concentration of 1000 cells per 50 µL.
4. **Seed aggregates**

- Add 50 µL of the cell suspension to each well of a 96-well U-bottom ultra-low attachment plate (ULU-96) (e.g., faCelliate BIOFLOAT, Cat. No.: F202003).
- Place the plate in a humidified incubator at 37°C, 5% CO₂.

**<u>Day 1: One day after aggregation</u>**

1. **Microscopic observation**

- Examine each well under a microscope to confirm a single, round aggregate with smooth edges. Target diameter is ∼200 µm; if aggregates are ∼240 µm, proceed directly to Day 2.
- If the aggregates do not have the optimal size, adjust the seeding density on Day 0.

**<u>Day 2: Mesoderm induction</u>**

1. **Microscopic observation**

- Aggregates should remain round with smooth edges and grow to a diameter of approximately 240 µm.
2. Prepare Mesoderm Induction Medium (8 mL total is sufficient for 96 wells)

- Thaw supplements to RT (21-24 °C).
- STEMdiff™ BVO Induction Basal Medium (STEMCELL, Cat. No.: 100-0654): 7.760 mL
- STEMdiff™ BVO Induction Supplement (STEMCELL, Cat. No.: 100-0655): 80 µL
- STEMdiff™ BVO Mesodermal Induction Supplement (STEMCELL, Cat. No.: 100-0656): 160 µL
- After mixing at RT, warm the Mesoderm Induction Medium to 37 °C.
3. Medium exchange

- Carefully aspirate all Aggregation Formation Medium from each well and replace with 70 µL of freshly prepared Mesoderm Induction Medium.
- Immediately return the plate to the incubator (37 °C, 5% CO₂).

**<u>Day 5: Vascular induction</u>**

1. **Prepare Vascular Induction Medium (8 mL total is sufficient for 96 wells)**

- Thaw supplements to RT (21-24 °C).
- STEMdiff™ BVO Induction Basal Medium (STEMCELL, Cat. No.: 100-0654): 7.760 mL
- STEMdiff™ BVO Induction Supplement (STEMCELL, Cat. No.: 100-0655): 80 µL
- STEMdiff™ BVO Vascular Induction Supplement (STEMCELL, Cat. No.: 100-0657): 160 µL
- After mixing at RT, warm the Vascular Induction Medium to 37 °C.
2. Medium exchange

- Carefully aspirate all Mesoderm Induction Medium from each well and replace with 70 µL of freshly prepared Vascular Induction Medium.
- Immediately return the plate to the incubator (37 °C, 5% CO₂).

**<u>Day 7: Embedding and blood vessel network formation</u>**

**1. Thaw reagents**

- Thaw STEMdiff™ Blood Vessel Organoid Maturation Medium (STEMCELL, Cat. No.: 100-0658) at RT (21-24 °C).

**2. Microscopic observation**

- Aggregates should appear round with flaky or rough edges and grow to a diameter of approximately 470 µm.

**3. Prepare nutrient mix on ice**

- The calculations below are given per 1 mL (nutrient mix plus collagen mix). However, prepare at least 5 mL total, as smaller volumes affect polymerization.
- Prepare the nutrient mix as follows per mL (adjust to needed volume):
- `o` Hams F12 Nutrient Mix (Gibco, Cat. No.: 11765-054): 159 μL
- `o` 5× DMEM/High Glucose (Cytiva, Cat. No.: SH30003.03): 125.2 μL
- `o` Sterile filtered water: 12.2 μL
- `o` GlutaMAX (Gibco, Cat. No.: 13462629): 6.4 μL
- `o` HEPES (Gibco, Cat. No.: 15630080): 12.6 μL
- `o` Sodium Bicarbonate (Gibco, Cat. No.: 25080-094): 9.8 μL

**4. Prepare desired extracellular matrix (ECM):**

- **Matrigel/Geltrex-bovine collagen mix:**
- `o` Use 750 μL collagen mix (PureCol, 3.2 mg/mL, Advanced BioMatrix, Cat. No.: m5005-100ML) + 250 μL Geltrex (Gibco, Cat. No.: A14133-02) or 250 μL Matrigel (Corning, Cat. No.: 356231).
- collagen (PureCol) (2.15 mg/mL):
- `o` Nutrient mix: 325.2 μL
- `o` Collagen (PureCol, 3.2 mg/mL, Advanced BioMatrix, Cat. No.: m5005-100ML): 666.6 μL
- `o` Sodium hydroxide (Sigma, Cat. No.: S2770-100ML): 8.34 μL
- collagen (2mg/mL):
- `o` Nutrient mix: 325.2μL
- `o` Collagen (3mg/mL) (CellAdhere™ Type I Collagen, Human, STEMCELL, Cat. No.: 07005) 650.4μL
- **NOTE:** The pH of the gel should be between 7.4 and 7.6.
- **NOTE:** Collagen concentration in Geltrex varies (∼2-5 mg/mL) according to the provider (Gibco) and Matrigel ∼2-6 mg/mL (Corning) respectively
- **NOTE:** For human collagen we found that collagen concentrations in the range of 1-5 mg/mL provided consistent polymerization and supported stable organoid morphology (see Supp. Fig. 3 A-C).
- Keep the extracellular matrix on ice to prevent polymerization.

**5. Embed aggregates**

- Completely remove the Vascular Induction Medium without disturbing the aggregates.
- Add 35uL of extracellular matrix (ECM) to the aggregates in the same ULU-96 plate (20-75 µL of ECM provides stable organoid formation, see Suppl. Fig. 1 H; Suppl. Fig. 2 A, B). The resulting culture of aggregate in ECM is termed “sitting drop”.
- For the sitting drops just let the appropriate amount of ECM drop on the media-free aggregate.
- The aggregate should be positioned in the center of the drop (not at the edges when viewed from above), and the drop should sit in the center of the well (not on the walls).
- Aggregates can be embedded in a collagen I-Matrigel (or Geltrex) mix (3:1) in collagen alone (e.g. PureCol) or in recombinant human collagen I.
- Incubate at 37 °C for at least 2 hours to allow for polymerization.

**6. Add Maturation Medium**

- After polymerization, add 100-150 μL of warm (37°C) STEMdiff™ BVO Maturation Medium (STEMCELL, Cat. No.: 100-0658) Maturation Medium without disturbing the sitting drops.
- Return the plate to the incubator (37 °C, 5% CO₂) immediately.

**<u>Day 8 (embedding day 1):</u>**

- Aggregates begin to differentiate into vascular networks. Examine the vascular networks under a microscope and look for radial sprouting.

**<u>Day 12 (embedding day 5)</u>**

1. **Microscopic examination**

- Observe the vascular networks / early BVOs under the microscope.
- Note that the gel may appear reduced in volume and partially pulled toward the BVOs. Some BVOs may begin to float in the medium.
2. Medium exchange

- Carefully aspirate the old Maturation Medium from each well, avoiding disruption to the BVOs or gel.
- Add 100-150 μL of fresh, pre-warmed (37°C) STEMdiff™ BVO Maturation Medium (STEMCELL, Cat. No.: 100-0658) Maturation Medium to each well.
- Immediately return the plate to a humidified incubator set at 37°C with 5% CO₂ to ensure optimal growth conditions.

**<u>Day 16-23: mature BVOs</u>**

1. **Microscopic examination**

1. BVOs should appear rounded, with residual gel at the edges or floating in the medium and are ready to use.
2. Extended culture

1. From day 16 onward, change the medium every 2–3 days. By day 23, BVOs should be free of gel.

### 2.3. Flow Cytometry

All data were acquired using a CytoFLEX S flow cytometer (Beckman Coulter) or a NAVIOS flow cytometer (Beckman Coulter) and analyzed with FlowJo software (version 10.10.0, Becton Dickinson). Briefly, three blood BVOs per sample were washed once with RPMI-1640 medium (Gibco, Cat. No.: 11875-093) and digested for approximately 40 minutes at 37°C using the Multi Tissue Dissociation Kit (Miltenyi Biotec, Cat. No.: 130-110-201). The digestion process was stopped by adding cold (4°C) RPMI-1640 supplemented with 10% fetal calf serum (FCS, Fisher Scientific, Cat. No.: 11573397). The cells were then centrifuged at 300 x g for 3 minutes, followed by a single wash with FACS buffer (composed of 1:20 BSA dilution using MACS® BSA Stock Solution, Miltenyi Biotec, Cat. No.: 130-091-376, in autoMACS® Rinsing Solution, Miltenyi Biotec, Cat. No.: 130-091-222). Subsequently, the cells were stained with the following antibodies: Anti-human CD31 (PE, REAfinity™, Clone: REA730, 1:100 dilution, Miltenyi Biotec, Cat. No.: 130-110-669, Lot: 5241009173); Anti-human CD140b (PDGFRβ) (REAfinity™, Clone: REA363, 1:100 dilution, Miltenyi Biotec, Cat. No.: 130-129-294, Lot: 5230704827); Anti-human TRA-1-60 (PE, REAfinity™, Clone: REA157, 1:100 dilution, Milteny Biotec, Cat. No.: 130-122-921, Lot: 5230401683; Anti-human SSEA-4 (APC, REAfinity™, Clone: REA101, 1:100 dilution, Milteny Biotec, Cat. No.: 130-123-815, Lot: 5230502976); Viability dye (eFluor™ 450, 1:1000 dilution, Invitrogen, Cat. No.: 65-0863-14, Lot: 2804457). Staining was performed for 30-60 minutes at RT in the dark. After staining, the cells were washed three times with FACS buffer and resuspended in 300-500 µL of FACS buffer for flow cytometric analysis (Gating strategy see Suppl. Fig 4).

### 2.4. Immunofluorescence Staining

Blood vessel organoids were transferred to 2 mL Eppendorf tubes using a cut 1 mL pipette tip. After settling, organoids were washed twice with D-PBS (Gibco, Cat. No.: 14190-094) and fixed in 2 mL of 4% paraformaldehyde solution (Fisher Scientific, Cat. No.: 28908) for 60 minutes at RT (21-24°C) on a rocking platform. Fixed organoids were washed twice with D-PBS and stored in D-PBS at 2-8°C for up to one month.

For blocking, organoids were incubated with 1 mL of blocking buffer [1% BSA (Roche, Cat. No.: 10735078001), 3% donkey serum (Biowest, Cat. No.: S2170-100), 0.5% Tween-20 (Sigma, Cat. No.: P1379-25ml), 0.5% Triton X-100 (Sigma, Cat. No.: X-100-100ml), 0.01% sodium deoxycholate (Sigma, Cat. No.: D6750-10G, from 1% wt/vol stock solution), and 94.99% D-PBS] at RT for 3 hours on a rocking platform.

Primary antibodies, including mouse anti-human CD31 (Agilent, Cat. No.: M0823, Lot: 41355845, Clone: JC70A, 1:100) and rabbit anti-human PDGFRβ (Cell Signaling, Cat. No.: 3169S, Lot: 15, Clone: 28E1, 1:100), were diluted in blocking buffer. A total of 100 µL of the primary antibody mix was added to the organoids, and the samples were incubated at 4°C overnight on a rocking platform. Following incubation, organoids were washed three times with wash buffer (0.05% Tween-20 in D-PBS), with a 30-minute rocking platform incubation during the third wash.

Secondary antibodies, including Alexa Fluor™ 546 donkey anti-mouse IgG (H+L) (Invitrogen, Cat. No.: A10036, Lot: 2306765, 1:250) and Alexa Fluor™ 647 donkey anti-rabbit IgG (H+L) (Invitrogen, Cat. No.: A31573, Lot: 2752586, 1:250), were diluted in blocking buffer. A total of 250 µL of the secondary antibody mix was added to the organoids, followed by a 3-hour incubation at RT in the dark on a rocking platform. Organoids were washed twice with wash buffer, leaving the second wash for 30 minutes in the dark on a rocking platform.

For clearing, organoids were incubated with 200 µL RapiClear 1.49 (SUNJin Lab, Cat. No.: RC149002) overnight. After clearing, organoids were transferred onto a coverslip with a 1 mm iSpacer (SUNJin Lab, Cat. No.: IS012), and residual RapiClear was carefully removed. For mounting, 300-400 µL RapiClear was applied before placing a coverslip on top. Coverslips were sealed with clear nail polish and stored at 4 °C overnight.

Image acquisition was performed using a confocal microscope (ZEISS Cell Observer SD), equipped with Neofluar 10x NA 0.3 Objective and Zen 2.6 software (ZEISS, blue edition, Version 2.6.76.00000). Image processing was carried out with the open-source software ImageJ2 (Version 2.16.0/1.54p).

### 2.5. Mouse Experiments

Five 8-week-old, male, NOD SCID gamma mice (NSG®: NOD.Cg-PrkdcSCID Il2rgtm1Wjl/SzJ, Charles River Laboratories, l’Arbresle, France) were individually housed in cages containing a sizzle nest, square pieces of paper, and a dome for environmental enrichment. The mice were maintained on an inverted 12-hour light/12-hour dark cycle (lights off from 8:00 AM to 8:00 PM), with a RT of 24±2° C and relative humidity of 50±20%. On Day 0 of the study, the back hair of each mouse was shaved under gas anesthesia with isoflurane (Iso-Vet, Piramal Critical Care).

A full-thickness, 12-mm diameter circular skin wound, including the panniculus carnosus, was then induced on the back of each mouse using a sterile biopsy punch (Acuderm Inc.). A donut-shaped silicone ring, 1-mm thick, 16-mm inner diameter and 24-mm outer diameter, prepared with transparent silicone sheets (Folioxane® Unrestricted, 0.5 x 90 x 150 mm, Novatech SA, Product no. FU050M, Batch no. 32226), and a 16 and 24 mm diameter punch system (Facom, Morangis, France) designed specifically, was glued around the wounded region, with special silicone glue (Kwik-Sil Silicone Elastomer low viscosity glue, World Precision Instrument) to prevent wound contraction. Subsequently, 10 BVOs were implanted into the wound bed. An Optiskin (Laboratoires URGO, Chenôve, France, Product no.OPTISKIN FILM 10cmx12cm, Batch no. 191205B) cover dressing and plaster strips were applied to secure the wound and silicone ring on all mice.

Mice were weighed twice per week at each renewal of the cover dressings and plasters. Daily observations of the mice were conducted to monitor mortality, behavior, and general condition. Upon complete healing of the skin wound, mice were euthanized via an overdose of pentobarbital. The harvested tissues were fixed in 4% PFA for 24 hours, then transferred and stored in 1X PBS.

All mouse experiments were conducted by ETAP-Lab (Nancy, France). The protocol received project authorization for animal use in scientific purposes from the Ministry of Higher Education and Research (No. APAFIS#51374) following ethical approval submission.

### 2.6. Statistics

Statistical analyses were conducted using GraphPad Prism. Details of the statistical tests applied are provided in the figure legends. A p-value of less than 0.05 was considered statistically significant.

## 3. Results

### 3.1. Aggregation in 96 well Plates and Implementation of the “Sitting Drops” Method for ECM Embedding

Prior to initiating blood vessel organoid (BVO) formation, we first confirmed the pluripotency of the human induced pluripotent stem cells (hiPSCs) used in our experiments by assessing the expression of TRA 1-60 and SSEA-4 (Supp. Fig. 1A). We then aggregated hiPSCs in ultra-low attachment 6-well plates according to the standard protocol by Wimmer et al. (6). After seven days (after mesoderm and vascular induction), this approach resulted in multiple aggregates of varying sizes (Fig. 1A), which strongly influenced downstream differentiation efficiency. These size discrepancies significantly affected the ratio of endothelial (CD31-positive) and pericyte (PDGFRβ-positive) cells in mature (day 16) BVOs (Fig. 1B). Notably, smaller aggregates contained higher proportion of CD31-positive cells than large ones, and PDGFRβ-positive pericytes were nearly absent in aggregates with diameter between 100 µm and 200 µm. Aggregates smaller than 100 µm failed to develop into mature BVOs altogether.

**Figure 1.**
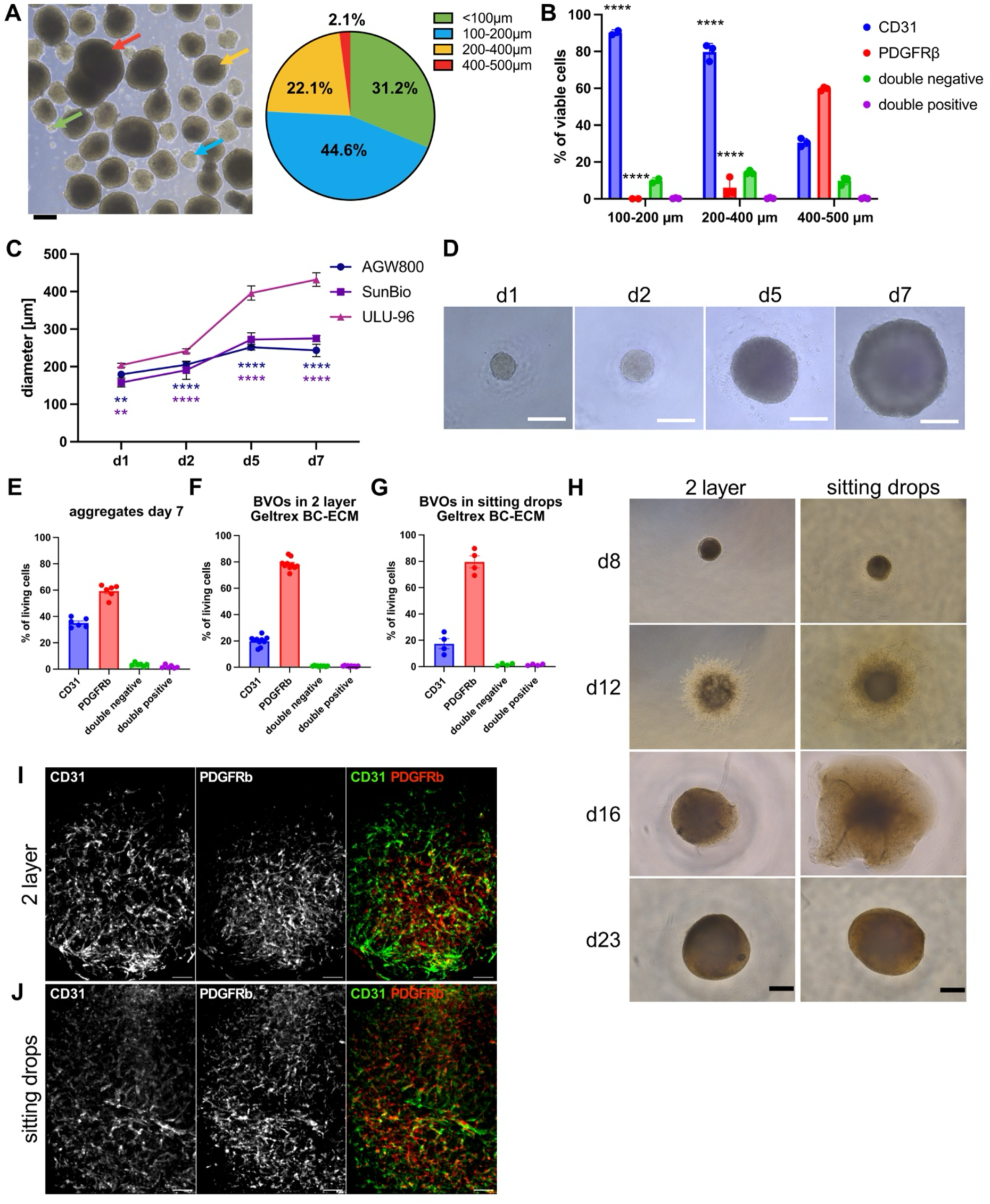
Optimization of blood vessel organoid (BVO) generation using ultra-low adhesion U-bottom 96-well plates. **A:** Brightfield microscopy image (10x) of human induced pluripotent stem cell (hiPSC) aggregates with size classification on day 6. Scale bar: 250 µm. Colored arrows indicate representative aggregate sizes corresponding to the categories shown in the pie chart: green (<100 µm), blue (100-200 µm), yellow (200-400 µm), red (400-500 µm). **B:** Flow cytometry analysis of BVOs on day 16 showing endothelial cells (CD31⁺) and pericytes (PDGFRβ⁺). Prior to embedding, aggregates were size-fractionated by sieving, and BVOs were generated from aggregates of defined diameter (100-200 µm, 200-400 µm, 400-500 µm). Aggregates with diameter smaller than 100 µm did not develop into BVOs and were excluded from the analysis. All statistical comparisons were performed relative to the 400-500 µm group. n = 3 technical replicates. Statistical analysis: two-way ANOVA with Tukey’s post hoc test. **C:** Comparison of aggregate growth in different culture plates: AggreWell 800 (AGW800), SunBio aggregation plate, and ultra-low attachment U-bottom 96-well plate (ULU-96). Aggregate diameters were measured on day 1 (post-aggregation), day 2 (prior to mesoderm induction), day 5 (prior to vascular induction), and on day 7 (prior to embedding). Data from two independent experiments (4-5 technical replicates each). Statistical analysis: two-way ANOVA with Tukey’s post hoc test. **D:** Brightfield images (10x) of developing aggregates in ULU-96 plates from day 1 to day 7 (same time points as in C). Scale bar: 200 µm. **E-G:** Flow cytometry analysis of CD31⁺ and PDGFRβ⁺ populations in (E) aggregates on day 7 (n = 6 biological replicates), (F) BVOs embedded in two layer Geltrex-bovine collagen mix ECM (BC-ECM) on day 16 (n = 4 biological replicates), and (G) BVOs cultured in Geltrex-BC-ECM as sitting drops in ULU-96 plates until day 23 (n = 4 biological replicates). **H:** Brightfield images (4×) showing development of BVOs embedded in Geltrex-(BC-ECM) as two-layer versus sitting drops cultures in ULU-96 plates on days 8, 12, 16, and 23. Scale bar: 500 µm. **I, J:** Immunofluorescence staining (10x) of BVOs on day 16. (I) BVOs were generated using the two-layer Geltrex-BC-ECM protocol or (J) the sitting drop method in ULU-96 plates with Geltrex-BC-ECM, and stained for CD31 (green) and PDGFRβ (red). Single-channel and merged images are shown. Scale bar: 100 µm. **Statistics:** **p < 0.01; ***p < 0.001.

To achieve uniform aggregate formation and improve differentiation consistency, we tested a range of commercially available plate types, some of them specifically designed for organoid culture. These included AggreWell 800 and SunBio plates, as well as various 96-well formats differing in well shape (U-bottom, flat-bottom) and surface coatings (ultra-low adhesion, delta surface, uncoated) (Fig. 1 D, Supp. Fig. 1B). Only the specialized aggregation plates (AggreWell 800 and SunBio plates) and U-bottom ultra-low attachment 96-well plates (ULU-96) generated single, compact aggregates per well after one day. However, only ULU-96 plates supported continuous and homogeneous growth during mesoderm and vascular induction (Fig. 1C). In contrast, aggregates in other formats either did not form properly, resulted in multiple irregular spheroids per well, or failed to grow consistently. Aggregates formed in ULU-96 plates maintained uniform size, spherical shape, and exerted stable morphology across all stages of differentiation (Fig. 1C, D).

Flow cytometric analysis further confirmed these observations. Aggregates cultured in ULU-96 showed markedly lower proportions of double-negative (CD31−/PDGFRβ−) cells (d7: 3.5±1.2%; d16: 1.6±0.9%) compared to those formed in AggreWell (d7: 82.1±1.8%; d16: 23.1±3.0%) or SunBio plates (d7: 75.0±8.0%; d16: 22.6±4.1%), indicating better differentiation efficiency (Fig. 1E, F; Supp. Fig. 1C-F). Other 96-well plate types produced irregular, poorly formed aggregates that lacked the size and structural integrity required for robust differentiation (Supp. Fig. 1B). Based on these findings, we selected ULU-96 plates as the standard format for aggregate formation moving forward.

In the conventional two-layer protocol by Wimmer et al. (6), vascular networks are embedded at day 7 between two layers of a Matrigel-bovine collagen mix ECM (BC-ECM) in 12-well plates. However, we observed substantial batch-to-batch variability in Matrigel, which led to inconsistent differentiation outcomes (Supp. Fig. 1G). To overcome this limitation, we established Geltrex as a standardized ECM substitute for Matrigel based on its reproducible performance. While CD31 and PDGFRβ expression levels varied significantly across different Matrigel lots, Geltrex consistently supported stable differentiation of both endothelial and pericyte lineages (Supp. Fig. 1G) and was therefore adopted as our extracellular matrix of choice. Importantly, immunofluorescence staining confirmed that BVOs generated using Geltrex exhibited well-organized vascular networks containing CD31 and PDGFRβ positive cells, closely resembling those obtained with Matrigel (Fig. 1I, Supp. Fig. 1I).

To optimize the protocol and eliminate the excision step typically performed on day 12, we tested whether ECM embedding could be carried out directly in the ULU-96 aggregation plates. On day 7, following vascular induction, we removed the medium and applied 35 µL of Geltrex-BC-ECM directly onto the aggregates, allowing continued culture in the same wells without transferring the BVOs. We refer to this simplified embedding approach as the “sitting drop” method. ECM volumes ranging from 20-75 µL resulted in CD31 and PDGFRβ expression profiles (Supp. Fig. 1H) comparable to those of BVOs embedded using the conventional two-layer method (Fig. 1F). The ECM volume was carefully chosen so that by day 12, matrix digestion and contraction allowed the BVOs to detach naturally, eliminating the need for mechanical excision. Immunofluorescence and brightfield imaging confirmed the development of spherical organoids with vascular structures closely resembling those generated by the standard embedding protocol (Fig. 1H-J, Supp. Fig. 2A, B).

**Figure 2.**
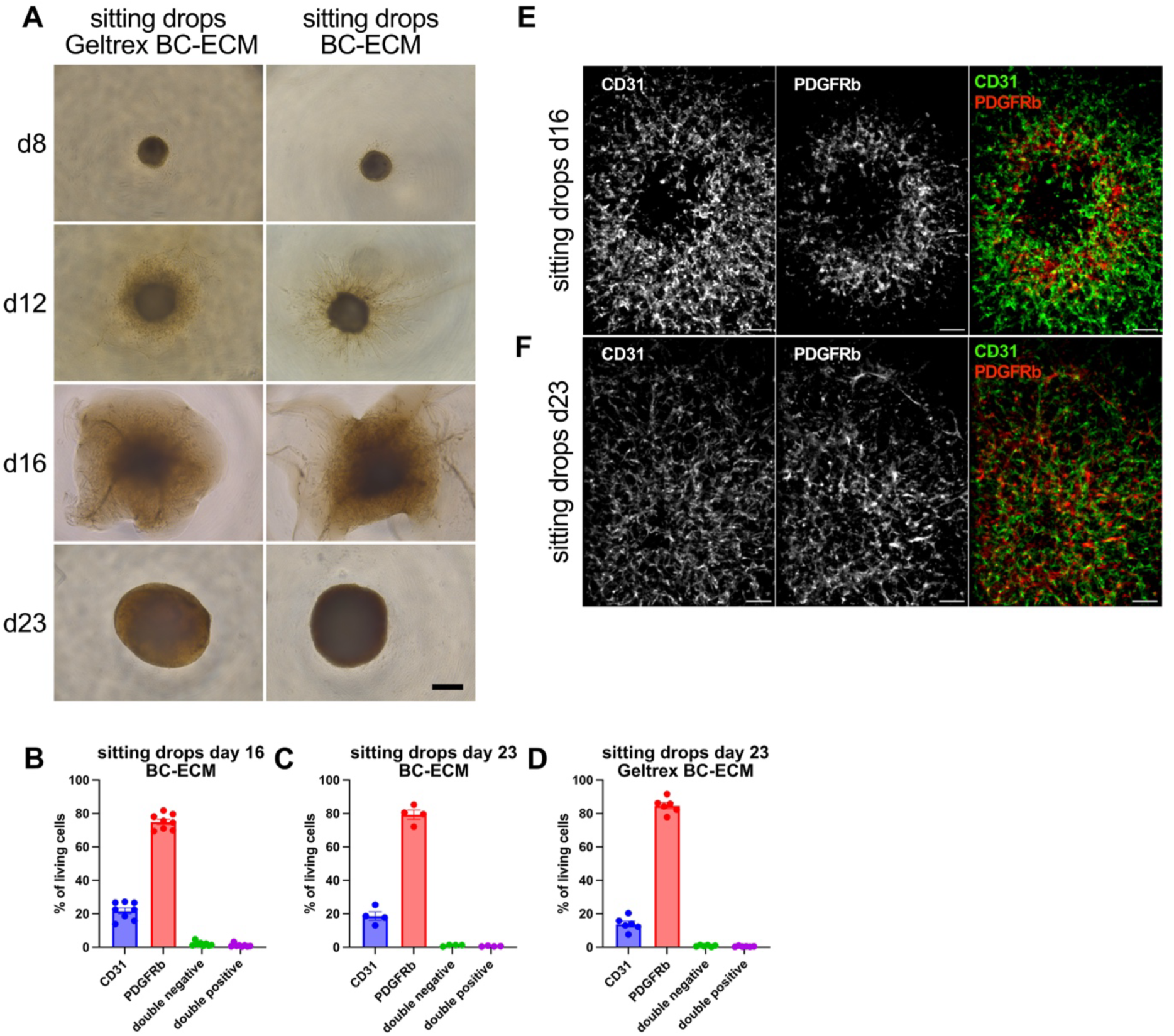
Generation of blood vessel organoids (BVOs) in a defined bovine collagen extracellular matrix (BC-ECM). **A:** Brightfield microscopy images (4x) comparing development of BVOs (sitting drops) embedded in bovine collagen (BC-ECM) versus Geltrex-BC-ECM. Images were taken on days 8, 12, 16, and 23. BVOs in both ECM types developed a similar spherical morphology over time. Scale bar: 500 µm. **B:** Flow cytometry analysis of BVOs (sitting drops) embedded in BC-ECM at day 16, showing proportions of endothelial cells (CD31⁺, blue), pericytes (PDGFRβ⁺, red), double-negative (green), and double-positive (purple) populations. n = 4 biological replicates with 2 technical replicates each. **C:** Flow cytometry analysis of BVOs (sitting drops) in BC-ECM at day 23. Cell composition remained comparable to day 16. n = 4 biological replicates. **D:** Flow cytometry analysis of BVOs (sitting drops) embedded in Geltrex-BC-ECM at day 23. n = 4 biological replicates. **E, F:** Immunofluorescence staining (10x) of BVOs generated as single-layer sitting drops in ULU-96 plates with BC-ECM, shown at day 16 (E) and day 23 (F). Samples were stained for CD31 (green) and PDGFRβ (red); single-channel and merged images are shown. Scale bar: 100 µm.

**Figure 3.**
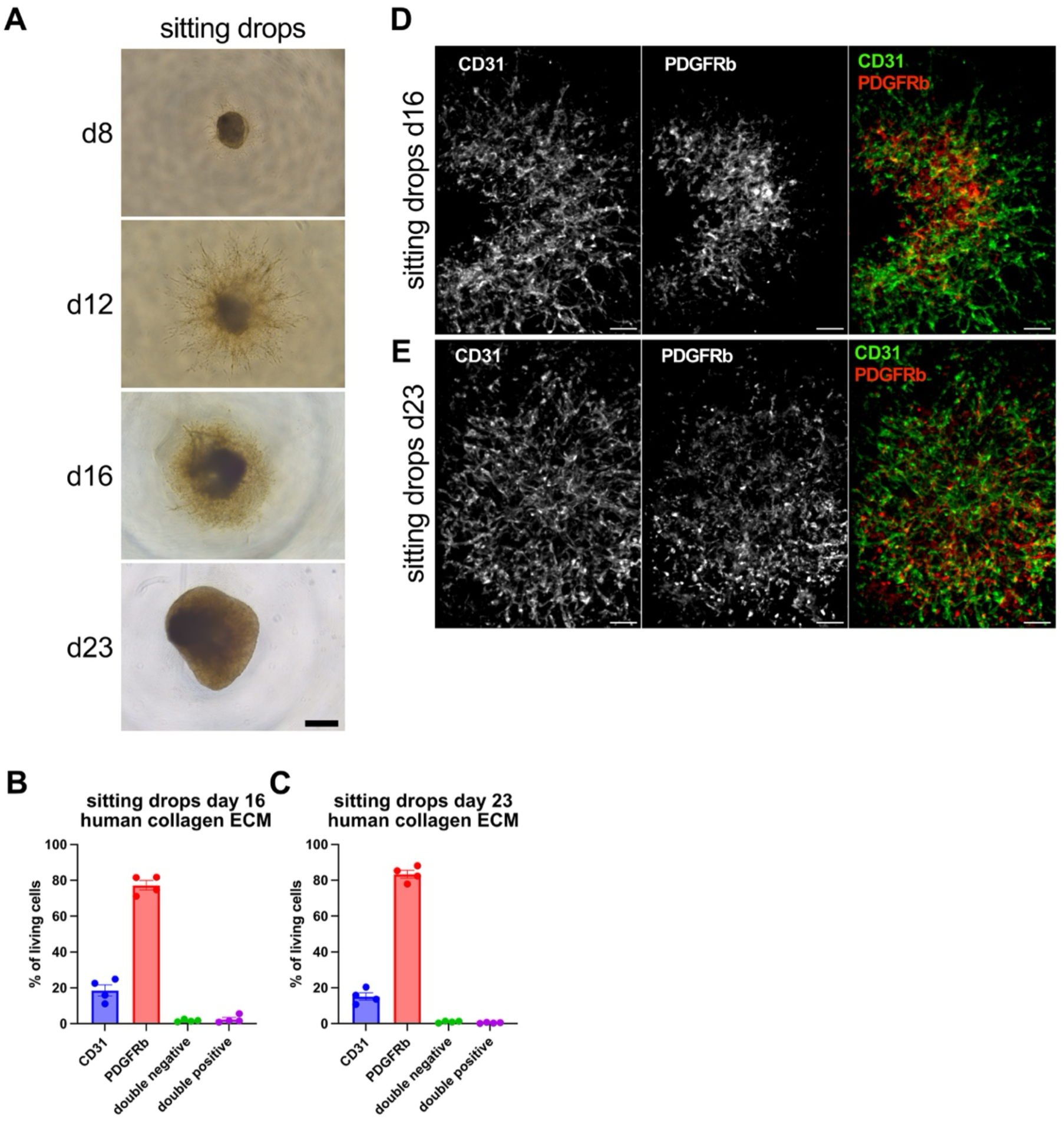
Generation of animal-origin-free blood vessel organoids (BVOs) using recombinant human collagen I. **A:** Brightfield microscopy images (4x) of BVOs embedded in human collagen ECM (HC-ECM) using the sitting drop method in ultra-low attachment U-bottom 96-well (ULA-96) plates. Images were taken on days 8, 12, 16, and 23. Scale bar: 500 µm. **B, C:** Flow cytometry analysis of BVOs embedded in HC-ECM using the sitting drop method at day 16 (B) and day 23 (C), showing proportions of endothelial cells (CD31⁺, blue), pericytes (PDGFRβ⁺, green), double-positive cells (red), and double-negative cells (purple). Cell composition remained stable over time. n = 4 biological replicates. **D, E:** Immunofluorescence staining (10x) of BVOs embedded in HC-ECM using the sitting drop method and cultured in ULU-96 plates, imaged at day 16 (D) and day 23 (E). Samples were stained for CD31 (green) and PDGFRβ (red); single-channel and merged images are displayed. Scale bar: 100 µm.

Low ECM volumes (:s20 µL) led to the generation of more compact organoids, whereas higher volumes (?:75 µL) resulted in loosely organized structures with reduced vessel network complexity and delayed spheroid maturation, extending up to day 30 (Supp. Fig. 2B).

We evaluated multiple 96-well plate types for ECM embedding of sitting drops and found that only ULU-96 plates consistently supported proper 3D development (Supp. Fig. 1J). By day 12, BVOs in alternative plate types developed dense, non-vascularized cores and failed to organize into vascular networks. A major advantage of ULU-96 plates is their compatibility with both aggregation and embedding steps, allowing the entire protocol to be executed in a single plate. This eliminates the need for manual transfers and significantly reduces variability and handling errors.

### 3.2. Generation of BVOs in a defined, Matrigel-/Geltrex-Free ECM

Matrigel and Geltrex are complex, animal-derived extracellular matrices that exhibit considerable batch-to-batch variability and have limited translational relevance (15, 16). To address these limitations, we established a method for generating BVOs using a defined ECM, free of both Matrigel and Geltrex. Aggregates were embedded in ULA-96 plates using 2.15 mg/mL final concentration of bovine-collagen (PureCol) as the sole source of collagen I, with 35 µL of ECM applied per sitting drop.

By day 16, BVOs generated in bovine-collagen ECM (BC-ECM) sitting drops displayed morphological features indistinguishable from those embedded in Geltrex-BC-ECM sitting drops, including a uniform spherical shape and homogeneous structure (Fig. 2A). Because the ECM in bovine-collagen cultures was not fully degraded by day 16, we extended the culturing period to day 23. At that point, the matrix was completely digested, and the BVOs had matured into compact, spherical structures (Fig. 2A, F).

Flow cytometry analysis revealed that the cellular composition of BVOs at day 23 grown in BC-ECM sitting drops closely resembled that of Geltrex-BC-ECM derived BVOs (Fig. 2C, D). Extending the culture from day 16 to day 23 had no impact on the cell composition in either condition (Fig. 2B-D, Fig. 1G), indicating high stability of mature BVOs. Morphologically, BVOs grown in BC-ECM at both time points were comparable to those generated in Geltrex-BC-ECM (Fig. 2E, F; Fig. 1I, J). Notably, immunofluorescence staining of BC-ECM sitting drops derived BVOs at day 23 revealed a slightly improved structural organization with more distinct vascular patterns (Fig. 2F). These findings demonstrate the feasibility of generating BVOs in a defined, Matrigel- and Geltrex-free ECM.

### 3.3. Generation of BVOs in Animal-origin-free ECM

To support the potential clinical application of BVOs in cell-based therapies and personalized medicine, it is essential to transition toward animal-origin-free culture conditions. Eliminating animal-derived components improves experimental reproducibility, minimizes the risk of contamination, and aligns with current regulatory and ethical standards. Moreover, animal-origin-free systems support safer, more standardized, and sustainable production processes.

To this end, we generated BVOs using commercially available recombinant human collagen type I as the component of the ECM. Aggregates were cultured in ULA-96 plates and embedded in 35 µL of 2 mg/mL final concentration of human collagen ECM (HC-ECM) per sitting drop (Fig. 3). Human collagen concentrations between 1 and 5 mg/mL yielded reproducible polymerization behavior and resulted in structurally stable BVOs (Supplemental Fig. 3). Flow cytometry analysis confirmed consistent formation of both endothelial (CD31⁺) and pericyte (PDGFRβ⁺) populations across all conditions (Suppl. Fig. 3A), with statistically significant but moderate differences, such as a lower proportion of CD31⁺ cells at 3.5 and 5 mg/mL compared to 2 mg/mL. Sitting drops BVOs generated in 1 mg/mL HC-ECM produced the smallest spheroids, whereas use of 2, 3.5, and 5 mg/mL resulted in progressively larger ones, with 5 mg/mL forming the largest organoids (Suppl. Fig. 3B). Immunofluorescence staining further confirmed the presence of CD31⁺ endothelial networks and PDGFRβ⁺ pericytes under all conditions (Supplemental Fig. 3C).

BVOs formed in human collagen (2 mg/mL) exhibited typical organoid architecture at both day 16 and day 23 (Fig. 3A, D, E) and maintained stable proportions of CD31⁺ endothelial and PDGFRβ⁺ pericyte populations over time, with no notable changes in cell composition between the two time points (Fig. 3B, C). This animal-origin-free approach enabled the generation of BVOs with a cellular composition and structural organization comparable to those obtained using BC-ECM or Geltrex-BC-ECM. Importantly, the use of human collagen type I also allows for the integration of GMP-compliant components in future applications.

### 3.4. Functional Testing of BVOs generated in sitting drops *in vivo*

To evaluate the functionality of BVOs generated in BC-ECM (2.15 mg/mL PureCol) as sitting drops, we implanted them into full-thickness skin wounds of non-obese diabetic severe combined immunodeficiency (NOD SCID gamma) mice. After complete wound closure, skin samples were excised and subjected to immunohistochemistry for human CD31 to assess whether organoid-derived cells had integrated into host tissue.

We identified blood vessels containing human CD31-positive cells, indicating successful incorporation of the BVO-derived cells. Furthermore, these blood vessels contained erythrocytes in their lumen, providing evidence that the newly formed vasculature was functional and connected to the host’s circulatory system (Fig. 4).

**Figure 4.**
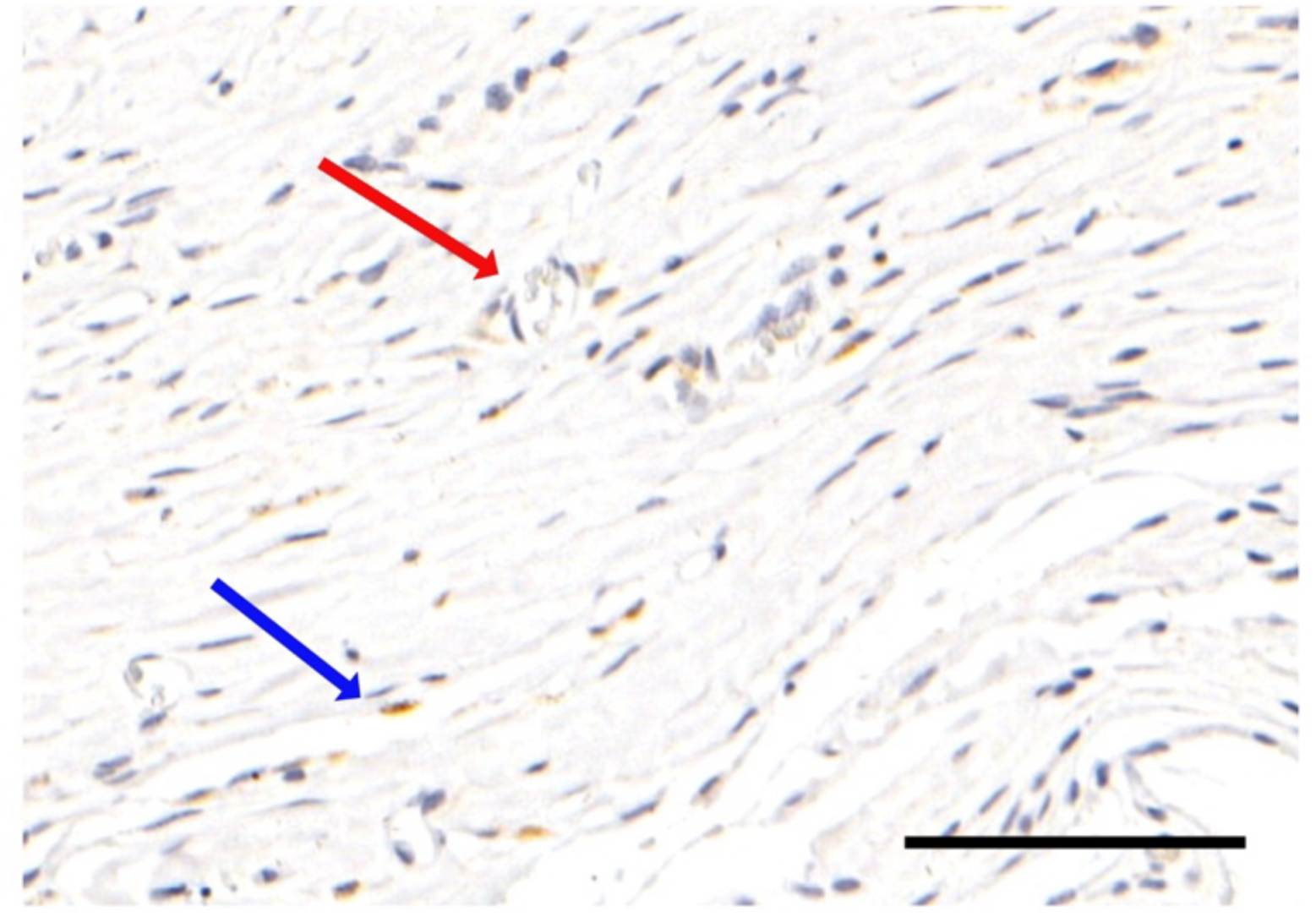
Immunohistochemistry of healed skin wounds, implanted with BVOs generated via the sitting drop method. Blood vessel organoids were implanted into open skin wounds, and tissue samples were collected after complete wound closure. Sections were stained with anti-human CD31 to identify vessel structures of organoid origin. The red arrow highlights a vessel in transverse section containing red blood cells, while the blue arrow indicates a vessel in longitudinal section. Scale bar: 100μm.

## 4. Discussion

The development of cell-based therapies represents a significant advancement in medical research. Organoid technologies, including patient-derived and allogeneic systems, are increasingly utilized for diagnostics, disease modeling, and as autologous grafts for regenerative therapies (11, 12). Blood vessel organoids (BVOs), in particular, offer a promising platform to support vascular repair, wound healing, and tissue regeneration due to their structural and functional resemblance to native vasculature (7).

Existing BVO production protocols are resource-intensive, laborious, and poorly suited for high-throughput applications (6, 7, 13, 14). A key limitation lies in the widespread use of animal-derived extracellular matrices such as Matrigel and Geltrex, which suffer from batch-to-batch variability, undefined composition, and lack of GMP compliance (15, 16). These properties restrict reproducibility and clinical applicability, posing regulatory and ethical challenges. Recent advances in organoid culture emphasize the need for defined, synthetic, or recombinant ECMs to overcome these challenges (17). Various animal-origin-free substrates can potentially provide viable support for pluripotent stem cell culture. A recent publication (18) identified Vitronectin as such promising animal-origin-free substrate for maintaining hiPSC pluripotency. In our experience, however, Vitronectin coatings resulted in premature differentiation, whereas Laminin-521 reliably preserved pluripotency over multiple passages and therefore provided a more stable starting point for downstream differentiation. The successful use of Fibronectin ECM in the same study (18), albeit more complicated and less viable for large scale utilization, outlines that there are further alternatives to animal-origin-free

ECM substrates. Notably, our findings align with the authors’ observations on the limitations of several commercially available hydrogel systems. Hydrogels from Manchester BIOGEL, GrowDex, VitroGel and Ectica did not support BVO formation in our hands, further underscoring the narrow range of biochemical and mechanical ECM properties compatible with vascular organoid differentiation.

In this study, we developed and validated an optimized and clinically relevant protocol for BVO generation using defined, animal-origin-free extracellular matrices (Fig. 5). By performing the entire process, from aggregation to organoid maturation, within a single ultra-low attachment 96-well plate, we reduced manual handling and contamination risk while enabling compatibility with robotic liquid handling platforms. This streamlined workflow allows for standardized large-scale production and facilitates the implementation of high-throughput drug testing. Compared to the widely used two-layer protocol by Wimmer et al. (6), our method eliminates excision steps, uses fewer reagents, and offers a simplified one-step ECM embedding procedure.

**Figure 5.**
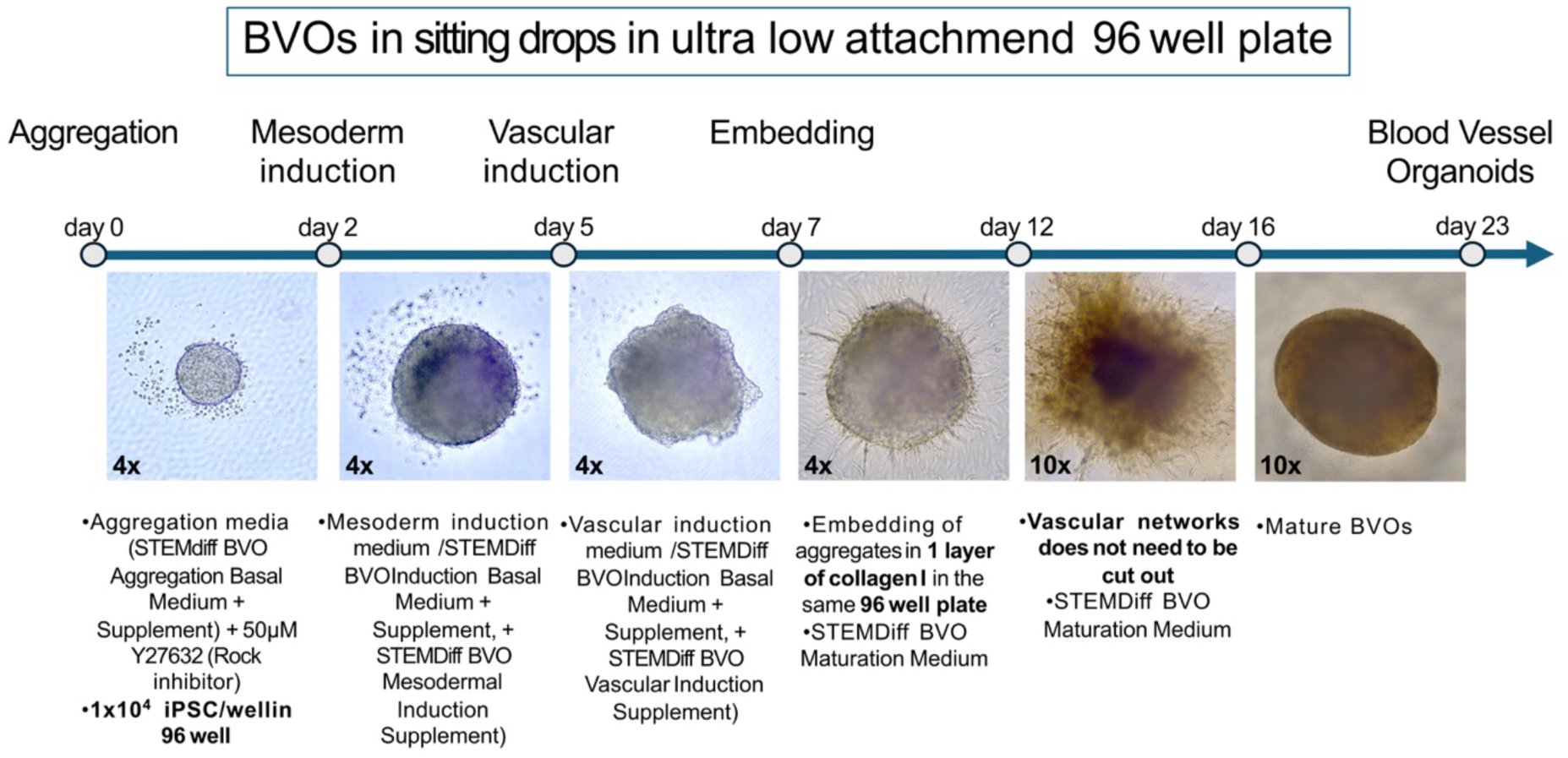
Schematic overview of the blood vessel organoid (BVO) generation protocol using the sitting drop method. Timeline representation of the BVO differentiation protocol performed in ultra-low attachment U-bottom 96-well plates. The scheme outlines each step of the process, including aggregation, mesoderm induction, vascular induction, and maturation, together with the corresponding reagents and media used at each time point. Representative brightfield images illustrate the progression of BVOs from day 1 (aggregation) to day 23 (mature organoid). The timeline highlights the standardized workflow and practical implementation for robust and reproducible BVO generation. Magnification is indicated in the bottom left corner of each image.

BVOs generated using this platform showed consistent morphology and a stable composition of CD31⁺ endothelial cells and PDGFRβ⁺ pericytes across different ECM types and culture durations. Immunofluorescence imaging confirmed comparable vascular organization between BVOs produced in Geltrex/Matrigel- and collagen-only-based matrices. Importantly, we demonstrate that recombinant human collagen type I is sufficient to support robust organoid formation, enabling a fully animal-origin-free workflow and GMP-compatible production. Across the tested concentration range (1–5 mg/mL), endothelial and pericyte differentiation remained stable, with statistically significant but moderate differences in CD31⁺ cell proportions at 3.5 and 5 mg/mL compared to 2 mg/mL. These findings highlight that the protocol is tolerant to collagen concentration variations within the ECM while maintaining reproducibility, which is critical for clinical translation.

*In vivo* implantation into full-thickness skin wounds in immunocompromised mice demonstrated functional integration of BVO-derived endothelial cells into host vasculature. While these results confirm integration with the host vascular network, future studies should investigate long-term stability, remodeling dynamics, and responses to ischemic or inflammatory stress. Moreover, evaluation in immunocompetent models will be necessary to investigate the interaction of BVOs with immune cells and potential rejection mechanisms.

Despite these advances, challenges remain. Polymerization behavior and mechanical properties of collagen-based ECMs can vary between lots and suppliers, potentially affecting reproducibility. Continued optimization and standardization of collagen formulations will be essential for scaling up production. Furthermore, integrating real-time monitoring and closed-loop feedback systems into automated workflows could enhance quality control for large-scale manufacturing (19).

While our attempts of using AggreWell or SunBio plates for the aggregation step resulted in suboptimal growth and an unfavorable CD31:PDGFRβ ratio in our hands, it is worth noting that other groups have reported successful aggregation and BVO generation using microwell platforms such as the AggreWell system (20, 21). Nevertheless, this did not limit our workflow, because the single ultra-low-attachment 96-well plate used here offers superior performance and significantly simplifies handling by eliminating transfers between plates.

Finally, emerging bioengineering technologies such as microfluidic platforms, 3D bioprinting and multi-organoid assembloids provide additional opportunities to enhance vascular architecture and complexity and integration (22–24). Combining our protocol with patient-specific iPSC lines could pave the way for individualized vascular organoids in disease modeling, drug screening, and ultimately autologous therapies.

In summary, we present a robust and efficient method for generating BVOs in a single-plate format using GMP-compatible and animal-origin-free components. The sitting drop technique simplifies the workflow, supports high-throughput scalability, and provides a foundation for clinical translation of vascular organoid technologies.

## 5. Conclusion

In summary, our study presents a novel, efficient, and robust protocol for the generation of blood vessel organoids. By eliminating animal-origin-derived components and simplifying the culture process, this approach offers a scalable and clinically relevant platform for organoid-based applications. Future work will focus on further optimizing the system for therapeutic use, including disease modeling, drug screening, and regenerative medicine.

## Author Contribution

A.H as first author designed and performed experiments, carried out the statistical analyses and wrote the manuscript. D.S. and J.T. designed and performed experiments and contributed to the writing of the manuscript. T.E.Y. initiated and supervised the project, designed experiments and wrote the manuscript as corresponding author.

## Data availability

The data, presented in this study, are available from the corresponding author on reasonable request.

---

**Supplemental Figure 1.**
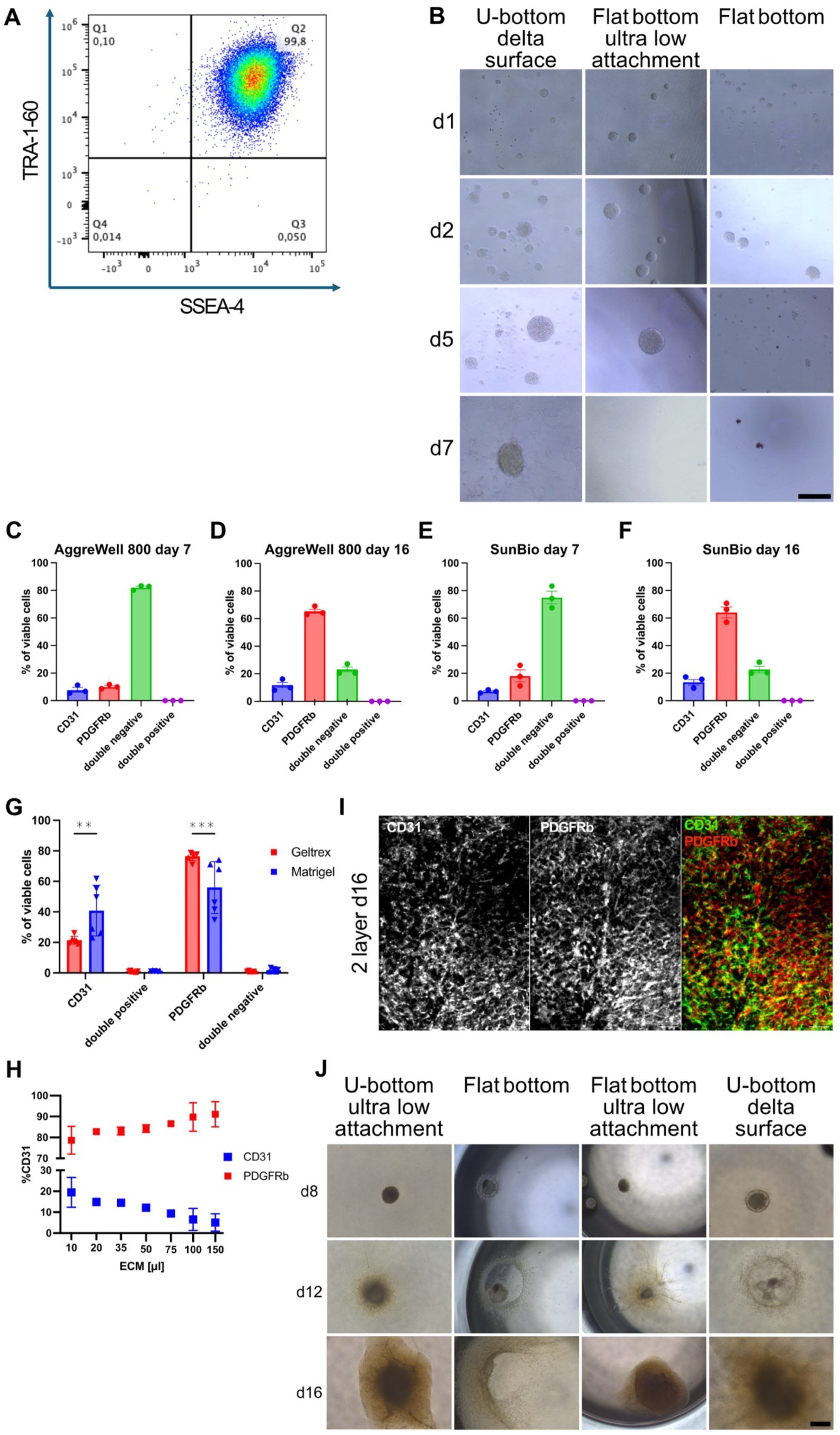
Supporting data for hiPSC pluripotency, plate testing, and BVO optimization. **A:** Flow cytometry analysis confirming pluripotency of hiPSCs prior to aggregation (day 0). Cells were stained for SSEA-4 and TRA-1-60. **B:** Brightfield microscopy images (10x) of cell aggregates cultured in different 96-well plate formats: delta surface U-bottom plates, flat-bottom ultra-low attachment plates, and standard flat-bottom plates. Images were taken on day 1 (post-aggregation), day 2 (prior to mesoderm induction), day 5 (prior to vascular induction), and day 7 (prior to embedding). Scale bar: 200 µm. **C-F:** Flow cytometry analysis of endothelial cells (CD31, blue), pericytes (PDGFRβ, red), double-positive cells (purple), and double-negative cells (green). (C) BVOs were generated using AggreWell 800 before embedding on day 7 and (D) after culture until day 16, or (E) using SunBio plates before embedding on day 7 and (F) after culture until day 16. N = 3 technical replicates per condition. **G:** Flow cytometry comparison of BVOs at day 16 generated with Geltrex-bovine collagen mix extracellular matrix (BC-ECM) (red) versus Matrigel-BC-ECM (blue). CD31⁺, PDGFRβ⁺, double-positive, and double-negative populations are shown. Data from 2 biological experiments with 3 technical replicates each. Statistical analysis: two-way ANOVA with Tukey’s post hoc test. **H:** Flow cytometry quantification of CD31⁺ and PDGFRβ⁺ cells in BVOs formed in ULU-96 plates, cultured with varying ECM volumes. BVOs were analyzed after reaching a spherical stage. Data represent 2 biological replicates. **I:** Immunofluorescence staining (10x) of BVOs on day 16, generated using the two-layer Matrigel-BC-ECM protocol and stained for CD31 (green) and PDGFRβ (red). Single-channel and merged images are shown. Scale bar: 100 µm. **J:** Brightfield microscopy images (4x) of BVOs embedded in 35 µL Geltrex-BC-ECM using the sitting drop method and cultured in ultra-low attachment U-bottom plates, standard flat-bottom plates, flat-bottom ultra-low attachment plates and delta-surface U-bottom plates. Images were taken on days 8, 12, and 16. Only BVOs in ULU-96 plates maintained a three-dimensional structure; in other plates, dense cores formed by day 12. Scale bar: 500 µm. **Statistics:** **p < 0.01; **\*\*\***p < 0.001.

**Supplemental Figure 2.**
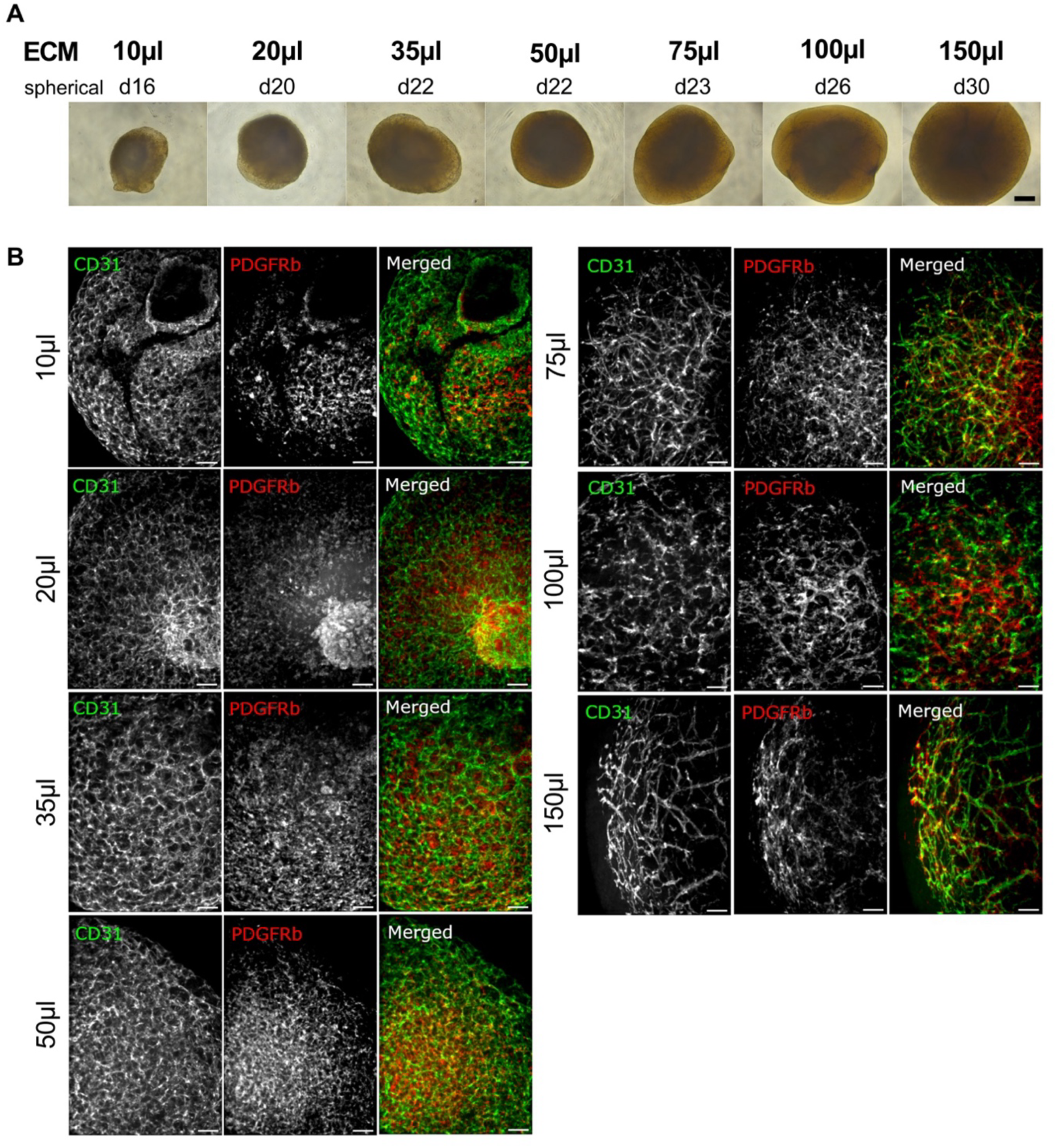
Development of BVOs using the sitting drop method in different volumes of extracellular matrix (ECM). **A:** Brightfield microscopy images (4x) of BVOs using the sitting drops method embedded in a Geltrex-bovine collagen mix ECM (BC-ECM). Sitting drops were cultured in different ECM volumes ranging from 10 µL to 150 µL. The day on which each BVO reached a spherical stage is indicated above the respective image. Scale bar: 500 µm. **B:** Immunofluorescence staining (10x) of BVOs using the sitting drops method after they got spherical, embedded in a Geltrex-BC-ECM. Sitting drops were cultured in different ECM volumes ranging from 10 µL to 150 µLand stained for CD31 (green) and for PDGFRβ (red). Single channels and merged images are shown. Scale bar: 100μm.

**Supplemental Figure 3.**
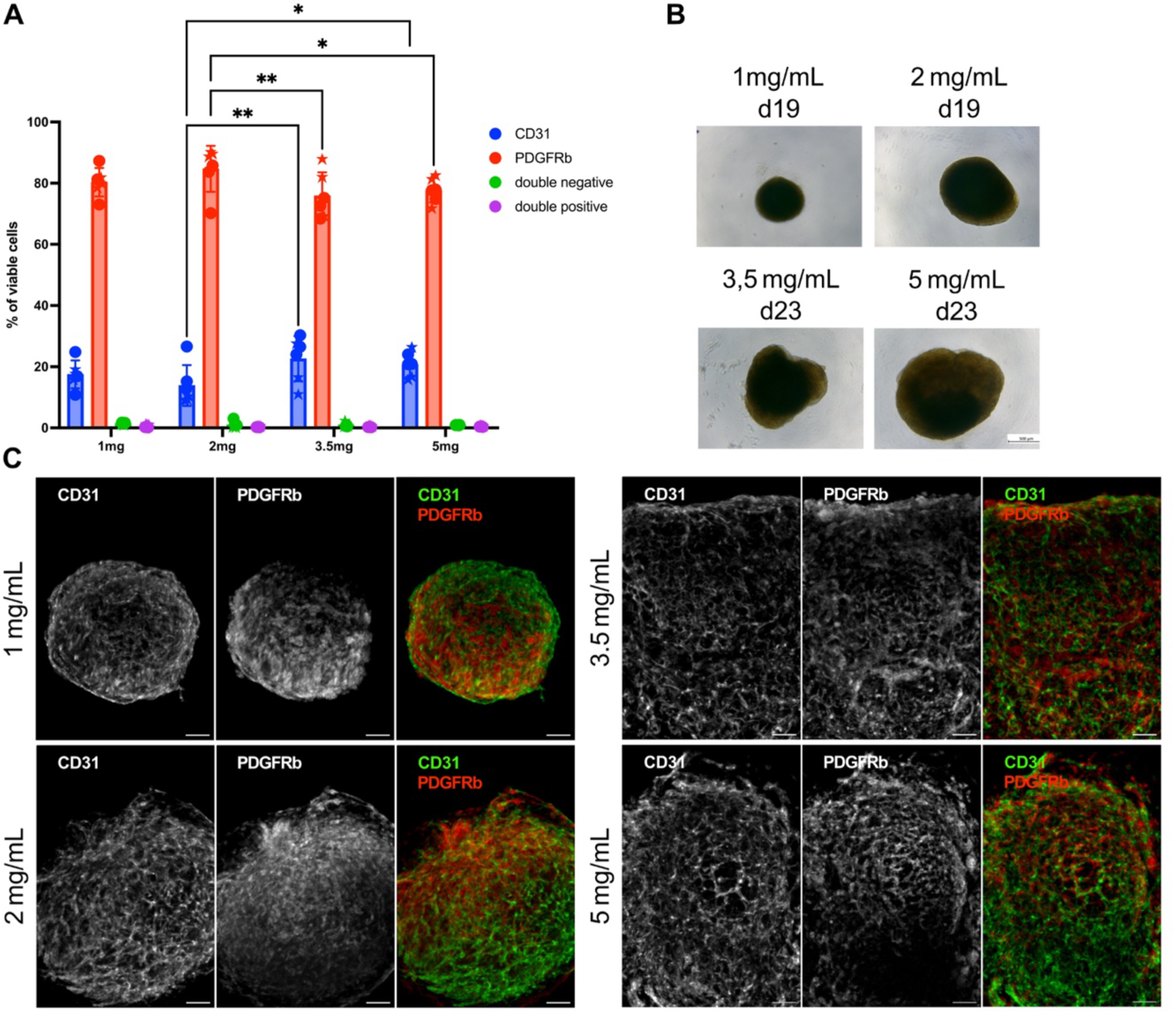
Development of blood vessel organoids (BVOs) using the sitting drop method in different concentrations of human collagen ECM (HC-ECM). **A**: Flow cytometry quantification of CD31⁺ and PDGFRβ⁺ cells in BVOs formed in ULU-96 plates using varying human collagen concentrations in the ECM. Sitting drops were cultured in different human collagen concentrations ranging from 1 mg/mL to 5mg/mL. BVOs were analyzed after reaching a spherical stage. Data represents 3 technical replicates. Statistical analysis: two-way ANOVA with Tukey’s post hoc test. **B**: Brightfield microscopy images (4x) of BVOs (sitting drops) embedded in HC-ECM. The day on which each BVO reached a spherical morphology is indicated above the respective image. Scale bar: 500 µm. **C:** Immunofluorescence staining (10x) of BVOs formed in ULU-96 plates using varying human collagen concentrations in the ECM upon reaching spherical appearance. BVOs were stained for CD31 (green) and for PDGFRβ (red). Single channels and merged images are shown. Scale bar: 100μm. **Statistics:** *p < 0.05**, p < 0.01.

**Supplemental Figure 4.**
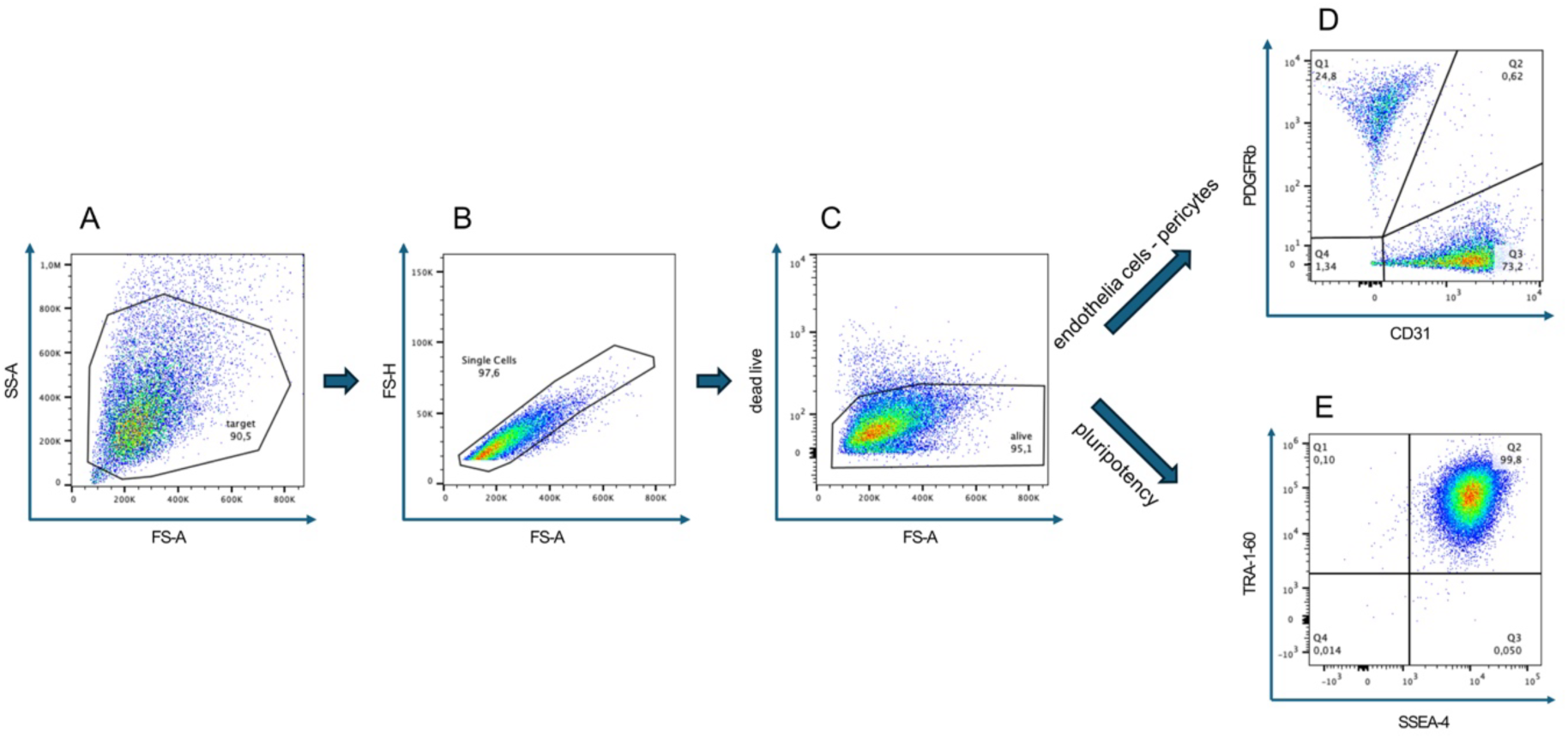
Flow cytometry gating strategy. **A-B:** Representative plots showing the gating strategy. (A) Debris, (B) doublets and (C) dead cells were excluded from the analysis. **D-E:** Viable cells from C were either analyzed for (D) endothelial (CD31) and pericyte (PDGFRβ) marker expression in vascular networks (day7) and mature organoids or (E) pluripotency (TRA-1-60 and SSEA-4) marker expression in induced pluripotent stem cells or at later stages of differentiation to confirm their absence.

